# Long-term, layer-specific reverberant activity in the mouse somatosensory cortex following sensory stimulation

**DOI:** 10.1101/058958

**Authors:** Elena Phoka, Aleksandra Berditchevskaia, Mauricio Barahona, Simon R Schultz

**Affiliations:** Department of Bioengineering, Imperial College London, SW7 2BP, London, UK; Department of Mathematics, Imperial College London, SW7 2RH, London, UK; Centre for Neurotechnology, Imperial College London, SW7 2BP, London, UK

**Keywords:** Barrel cortex, plasticity, spontaneous activity, electrophysiology

## Abstract

Neocortical circuits exhibit spontaneous neuronal activity whose functional relevance remains enigmatic. Several proposed functions assume that sensory experience can influence subsequent spontaneous activity. However, long-term alterations in spontaneous firing rates following sensory stimulation have not been reported until now. Here we show that multi-whisker, spatiotemporally rich stimulation of mouse vibrissae induces a laminar-specific, long-term increase of spontaneous activity in the somatosensory cortex. Such stimulation additionally produces stereotypical neural ensemble firing patterns from simultaneously recorded single neurons, which are maintained during spontaneous activity following stimulus offset. The increased neural activity and concomitant ensemble firing patterns are sustained for at least 25 minutes after stimulation, and specific to layers IV and Vb. In contrast, the same stimulation protocol applied to a single whisker fails to elicit this effect. Since layer Vb has the largest receptive fields and, together with layer IV, receives direct thalamic and lateral drive, the increase in firing activity could be the result of mechanisms involving the integration of spatiotemporal patterns across multiple whiskers. Our results provide direct evidence of modification of spontaneous cortical activity by sensory stimulation and could offer insight into the role of spatiotemporal integration in memory storage mechanisms for complex stimuli.

## Introduction

Even in the absence of sensory stimulation or motor output, neocortical circuits are not silent. Neurons exhibit highly stochastic *spontaneous* activity (Arieli et al., 1996;Azouz and Gray, 1999;Fox and Raichle, 2007;Ringach, 2009), which consumes a large fraction of the brain’s metabolic budget (Sokoloff et al., 1955). There is substantial evidence that structured spontaneous activity drives the assembly and refinement of sensory processing circuitry in the developing brain (reviewed in Kirkby et al, 2013). Although often considered to be ‘noise’ in the adult brain, spontaneous activity has a strong spatio-temporal structure: it reflects the functional architecture of cortical circuits (Tsodyks et al., 1999), resembles the patterns of activity produced by natural sensory stimulation, and propagates within cortical areas (Reyes-Puerta et al, 2016) and throughout brain-wide networks in reproducible patterns (Mitra and Raichle, 2016). These factors are suggestive of a role in information processing. In particular, spontaneous cortical activity has been demonstrated to affect sensory responses (Erchova et al., 2002;Ferezou et al., 2007;Kenet et al., 2003). It has also been suggested that spontaneous activity may reflect learning and memory processes (Lewis et al., 2009). However, despite substantial work, the basic mechanisms for such associations at the neural circuit level remain enigmatic.

For spontaneous activity to be a mediating factor in learning and memory, spontaneous neocortical firing rates and concomitant spatial patterns of activity should show long-term changes after sensory events (Tegnér et al., 2002). There have been indications, in several modalities, of transient changes due to sensory stimulation. For instance, extensive (24 hour) stimulation of mouse whiskers leads to an increase in inhibitory synaptic density in somatosensory cortex, and a transient reduction in spontaneous firing (Knott et al., 2002;Quairiaux et al., 2007). In prefrontal cortex, elevated firing rates are maintained for a short time in the absence of a sensory stimulus during delay-memory tasks (Goldman-Rakic, 1995), suggesting the possibility that information might be maintained during spontaneous firing by similar firing rate elevations. Increased correlation between sensory-evoked and subsequent spontaneous cortical activity (Han et al., 2008), as well as a reduction in response variability (Yao et al., 2007), has been found following stimulation. Although these experiments indirectly suggest a role for spontaneous activity in the maintenance of sensory information for learning and memory, there are currently no reports of sustained changes in firing rates following sensory stimulation or of maintained patterns of ensemble activity in subsequent spontaneous epochs following a purely sensory stimulus.

To address these issues, we performed electrophysiological recordings (both single- and multi-unit) from the anaesthetized mouse somatosensory cortex to test the effect on spontaneous activity of a naturalistic, spatio-temporally complex, multi-whisker stimulation protocol (Fig. 1*A*). Our results show that repetitive multi-whisker stimulation with a spatiotemporally structured stimulus leads to persistent changes in spontaneous firing rates. These changes are layer-specific (concentrated in layers IV and Vb) and long-lasting (beyond 30 minutes). Through mathematical analysis of our singleunit measurements, we also showed that particular ensemble firing patterns in these layers are maintained in the subsequent spontaneous activity. As multi-whisker stimulation involves the activation of trans-columnar afferent circuits, we hypothesized that if the effect involved strengthening of these connections, it would be absent when only one whisker was stimulated. We tested this in mice with all but the principal whisker trimmed, finding that indeed single-whisker stimulation does not induce the reverberant activity we observed with multi-whisker stimulation.

**Fig. 1.**
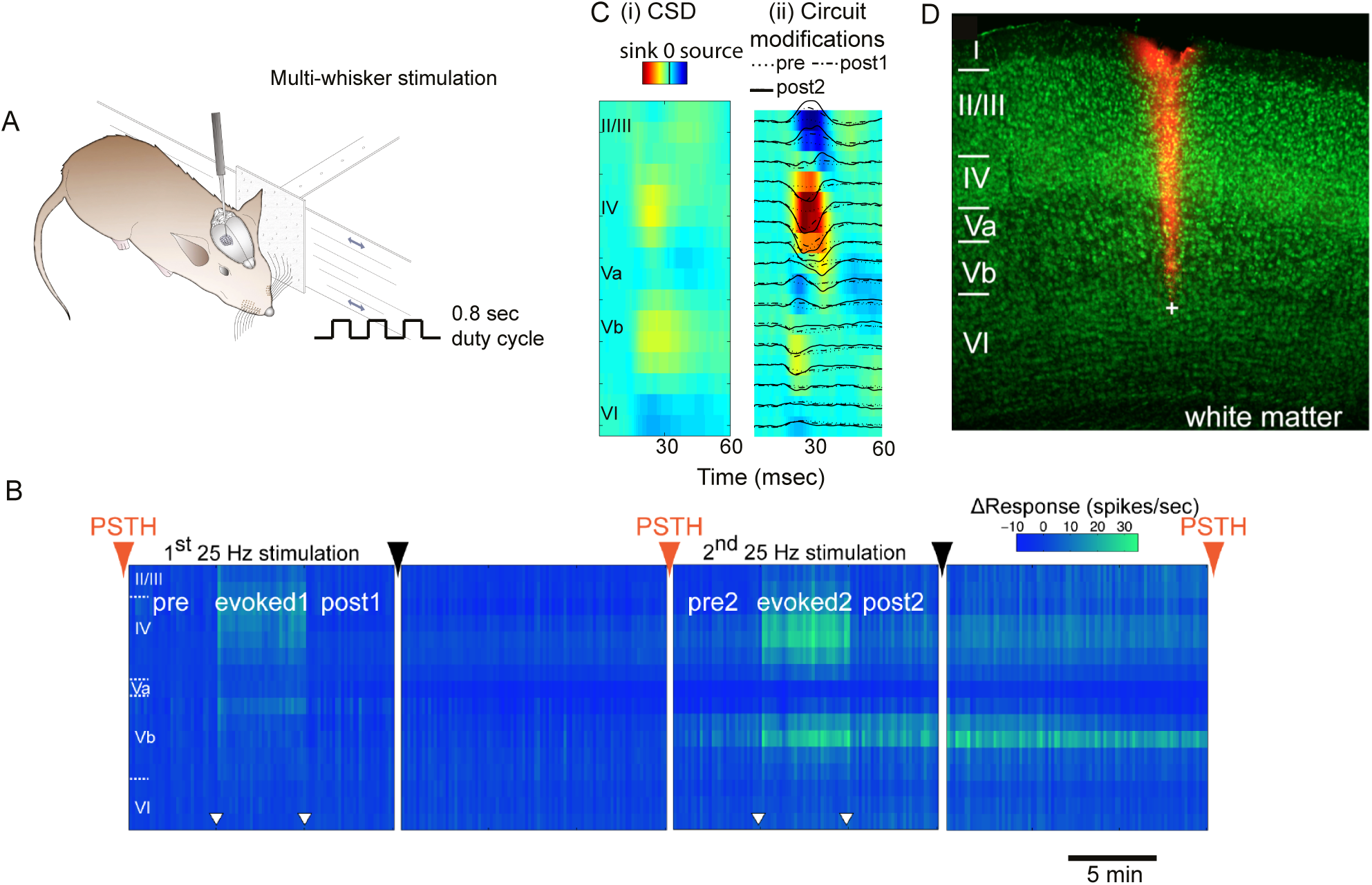
Long-lasting increase in spontaneous activity in layers IV and Vb following sensory stimulation. (*A*) Experimental setup: sandpaper swept across the mouse whiskers in the rostro-caudal axis while simultaneously recording from the contralateral somatosensory cortex. (*B*) Typical MUA recording from a linear probe spanning the cortical depth. Sensory stimulation at 25 Hz for 5 minutes (indicated within white triangles ▽) in two blocks induces an increase in spontaneous activity specific to layers IV and Vb, which is sustained for at least 25 minutes post-stimulation. 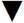 denotes a pause between the recordings; 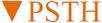 denotes brief stimulation trials designed to test sensory responsivity without inducing plasticity (see Fig. S1). (*C*) **(i)** Early sinks in layers IV and Vb (CSD, yellow) correspond to bands of increased spontaneous activity (CSD analysis of recording shown in panel B, from initial set of “PSTH” trials). **(ii)** Sensory stimulation induces a significant increase in the layer IV sink amplitude following the second stimulation (CSD analysis from final set of “PSTH” trials). (D) Layer boundaries were estimated by combining histological information (green channel, NeuroTrace fluorescent Nissl stain; red channel, DiI electrode coating) with CSD analysis.

## Results

### Long-lasting modification of neuronal firing rates in layers IV and Vb

To determine whether naturalistic sensory stimulation can modify the spontaneous firing rates of populations of neurons within a cerebral cortical circuit, we recorded spontaneous multi-unit neural activity (MUA) before and after collective stimulation of the facial whiskers of an anesthetized mouse with coarse sandpaper with an oscillatory motion at 25 Hz along the rostro-caudal axis (Fig. 1*A*, duty cycle 0.8 seconds ON and 0.8 seconds OFF, see Methods). We simultaneously recorded from several layers using a linear probe with 16 recording sites spanning the depth of the somatosensory cortex. We recorded spontaneous activity for 5 minutes (‘pre’), followed by 5 minutes of multi-whisker stimulation (‘evoked’), in turn followed by 5 minutes of post-stimulus spontaneous activity (‘post’), as shown in Fig. 1*B*. This protocol was performed twice in sequence.

Our experiments showed a striking increase in the post-stimulus spontaneous firing rates of layer IV and Vb neurons following the second stimulation (Fig. 1*B*). Laminar boundaries were established by histological analysis in direct correspondence with the two observed current source density (CSD) sinks of the barrel cortical column (Fig. 1*C,D*). The 16 electrode channels were assigned to cortical layers by careful alignment of histological identification of the probe tip (Blanche et al., 2005) with CSD analysis (Pettersen et al., 2006). The CSD distribution was calculated using the low-frequency component (1-300 Hz, LFP: local field potential) of the recorded signals (Mitzdorf, 1985;Freeman and Nicholson, 1975) and showed two current sinks appearing shortly after brief whisker stimulation at the locations of thalamocortical inputs (Swadlow et al., 2002), known to correspond to layers IV and Vb (Fig. 1*C,i*). We observed that the sink amplitude of the layer IV CSD distribution increased by a factor of 2.02±0.42 (ratio±s.e.m., P=0.019, N=7) following the first stimulation, and by a factor of 2.40±0.41 (P=0.005, N=7) following the second stimulation (see Fig. 1C, ii). The layer Vb CSD distribution remained relatively unchanged following the stimulation events (Fig. 1*C, ii*). The sink amplitude increase in layer IV, however, suggests the involvement of synaptic potentiation within and to layer IV (Mégevand et al., 2009).

Layer IV and Vb multi-unit sites exhibited an increase in spontaneous firing rates (post:pre ratio) directly related to the magnitude of the sensory response (evoked:pre ratio), which was strengthened with the repetition of the sensory events (Fig. 2). Some increase was also observed in layer VI, whereas layers II/III and Va showed no increase in spontaneous activity at all, regardless of the extent to which they were driven by sensory stimuli. The second stimulation epoch resulted in much more substantial changes in spontaneous firing rate (round symbols, Fig. 2B) for layers IV, Vb and VI in comparison to the first.

**Fig. 2.**
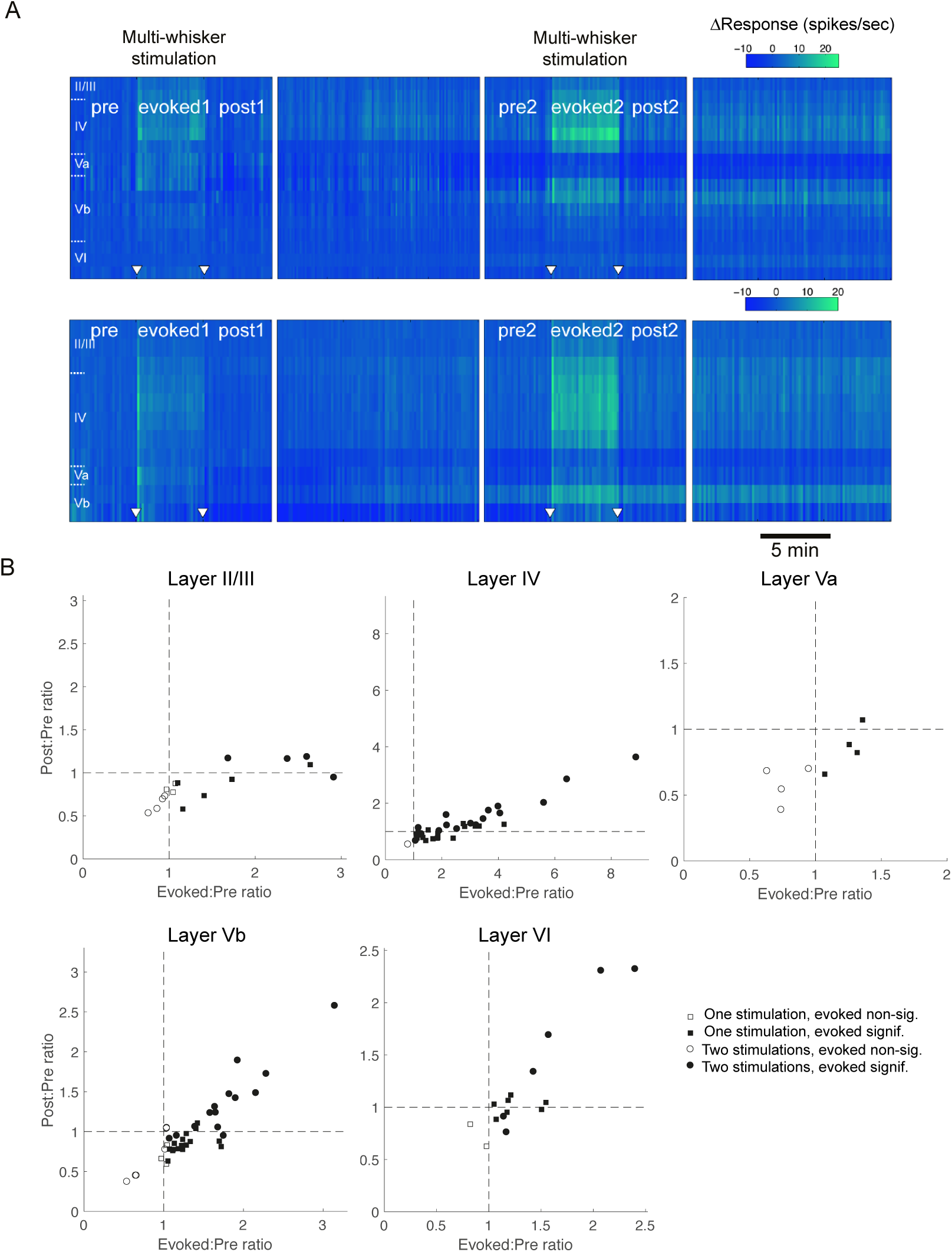
Modification of spontaneous activity following sensory stimulation (25 Hz) depends on the response during stimulation. (A) Two further examples of MUA laminar profiles, showing some initial, but more fragile, alteration of spontaneous activity after the first stimulation epoch, followed by more robust modification following the second stimulation epoch. (B) The spontaneous activity increase (Post:Pre ratio) is plotted against the response during stimulation (Evoked:Pre ratio) for MUA recorded in each of the channels in layers II/III, IV, Va, Vb and VI. Channels with significant responses *to the stimulation epoch* were identified by performing a one-tail (right) z-test to test the hypothesis that the evoked firing rate has a distribution significantly different to the distribution described by the mean and standard deviation of its pre-stimulus spontaneous firing rate. The significant channels so identified are indicated with filled symbols, and were used to calculate the population firing rates shown in Fig. 3. Activity following the first stimulation epoch is indicated by squares; activity following the second stimulation is indicated by circles.

Channels that exhibited a significant sensory response during the stimulation epoch, with significantly higher evoked firing rate than that of the pre-stimulus spontaneous activity (Fig. 2B, filled points), were identified and analyzed further (Fig. 3; bar chart data includes only data from channels showing statistically significant sensory-evoked activity during the stimulation epoch, i.e. filled points of Fig. 2B, to more specifically examine the consequences of that sensory response). Multi-unit sites with a significant sensory response in layer IV showed increased post-stimulus spontaneous activities by a ratio of 1.49±0.18 (p=0.0062, N=18) following the second stimulation (Fig. 3). In the case of layer Vb, sites with significant sensory responses (Fig. 2B, filled points) showed slightly decreased spontaneous firing rates (by a ratio of 0.85±0.03, P=6.9E-5, N=16, Fig. 3, empty asterisks) following the first stimulation, but increased their spontaneous ‘post’ activities by a ratio of 1.37±0.13 (P=0.0050, N=14, Fig. 3, filled asterisks) following the second stimulation. We conclude that while a single 5 minute stimulation epoch was sufficient to induce changes in spontaneous activity in some instances (see examples in Fig. 2A), the effect was not robust; in contrast, after repeating the stimulation epoch for a second time, a robust, layer-specific increase in spontaneous activity was observed.

**Fig. 3.**
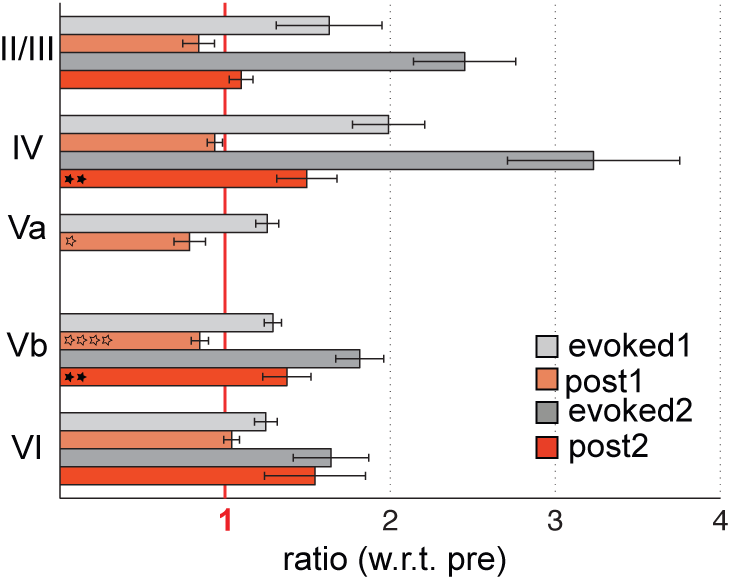
Spontaneous activity in layers IV and Vb spontaneous activity significantly decreases following repeated multi-whisker stimulation. Bar charts show mean (error bars s.e.m.) firing rates for the five minute colour-coded epochs, divided by the firing rate in the five minutes preceding the onset of stimulation, for sites showing statistically significant responses during the most recent *Evoked* epoch. Note that no electrode sites in layer Va remained significantly activated by the second sensory stimulation epoch (as apparent in the examples of Fig. 1 and Fig. 2A), and thus they are not considered here. Asterisks indicate significance of deviation from unity of the ratios from each such group of electrode sites across animals: ^*^p<0.05, ^**^p<0.01, ^***^p<0.001, ^****^p<0.00001.

The increase in spontaneous firing rates in layers IV and Vb (which showed the most robust effects) was long-lasting, with duration of at least 25 minutes post-stimulus (Fig. 4).

**Fig. 4.**
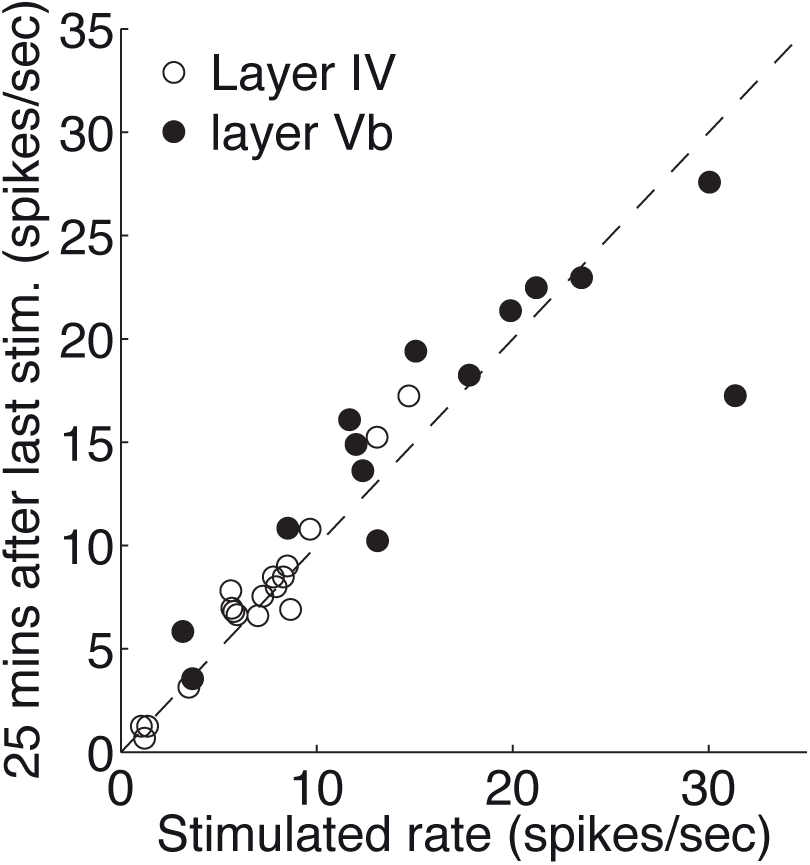
Increases in spontaneous firing rate in layers IV and Vb were sustained for at least 25 minutes. Spontaneous firing rate over a 30 second period immediately following the second stimulation (x axis), compared with the spontaneous rate over a 30 second period measured approximately 25 minutes later (y axis), for each of the channels in layers IV and Vb of the MUA. After 25 minutes have elapsed from the stimulation, 93% of the layer Vb MUA (13 of 14) and 94% of the layer IV MUA (17 of 18) channels remained within 10% of the firing rate recorded immediately after cessation of the stimulation.

### The increase in neuronal firing rates is unrelated to depth of anaesthesia

Sensory processing may be dynamically modulated by the intrinsic cortical state, depending on whether the cortical network is synchronized or desynchronized (Marguet and Harris, 2011;Goard and Dan, 2009), as well as by the level of anaesthesia (Chauvette et al., 2011). Transitions between such levels are accompanied by modifications in 1-4 Hz (Chauvette et al., 2011) or 1-8 Hz (Marguet and Harris, 2011;Goard and Dan, 2009) power of the LFP, and of neuronal firing rates (Harris and Thiele, 2011).

To examine the possibility that changes in depth of anaesthesia could play a role in our findings, we calculated a spectrogram using the LFP component of the recorded signal for each channel in layer Vb. We examined whether changes in power in the 1-4 Hz and 1-8 Hz frequency bands (post2:pre power ratio) relate to increases in the firing rates (post2:pre firing rate ratio). We found that the increase in firing rates is not accompanied by corresponding changes in power ratio in either 1-4 Hz or 1-8 Hz frequency bands for all sites within layer Vb and across experiments (Fig. 5 *B*(i) and *C*(i)). The mean values of LFP power in both bands showed no change between the prestimulus and post-stimulus spontaneous epochs: for 1-4 Hz, P=0.35 for a paired Student’s t-test, following the first stimulation and P=0.34 following the second stimulation; for 1-8 Hz, P=0.45 following the first stimulation and P=0.55 following the second stimulation. These results suggest that the overall cortical state and level of anaesthesia remained unchanged before and after the sensory events, and independent of modifications in the firing rates. We therefore consider it unlikely that the persistent firing rate increase of layer IV or Vb neurons is due to changes in depth of anaesthesia.

**Fig. 5.**
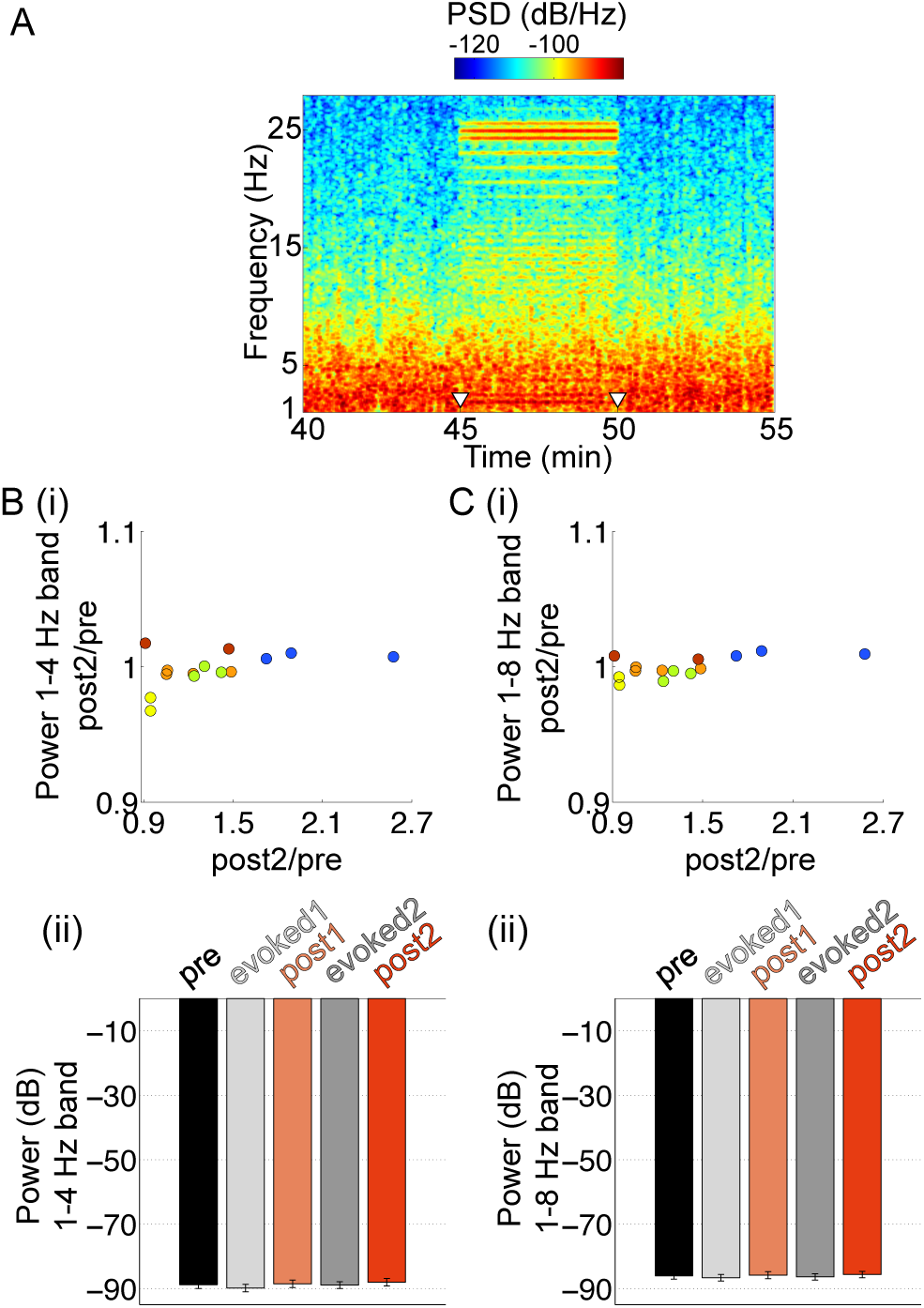
Spontaneous activity increase in layer Vb is not related to changes in depth of anaesthesia. (*A*) Typical spectrogram from a channel in layer Vb, recorded around the second 25-Hz stimulation epoch. The spontaneous spectral content pre- and post-stimulus is similar. Note the visible intermodulation, i.e. the frequency content modulation between the 25 Hz stimulus frequency and the 1.6 second 50% duty cycle. (*B*) (i) There is no significant change in power in the 1-4 Hz band from pre to post2 (ratio of power post2/pre), plotted against the change in firing rate (ratio post2/pre) for each individual recording site within layer Vb across experiments (N=14). Recording sites from the same animal have the same color-coding. (ii) Average power in the 1-4 Hz band decreases during stimulation (evoked1, evoked2) but returns to baseline in the spontaneous epochs (post1 and post2). Comparing post1 to pre and post2 to pre yields no statistically significant difference, indicating little change in the level of anaesthesia throughout the spontaneous recordings (N=14). (*C*) Same analysis as (*B*) for the 1-8 Hz band suggests that the cortical state more broadly remained at baseline throughout the spontaneous epochs during these experiments. Comparison of post1 to pre and post2 to pre in (ii) yielded no statistically significant differences.

### Single-whisker stimulation does not increase spontaneous activity in layers IV and Vb

Compared to neurons in the other cortical layers, layer Vb excitatory neurons have several distinct features: they receive excitatory inputs from all other cortical layers and neighbouring columns (Schubert et al., 2007b); they have larger receptive fields, as they respond to sensory inputs from approximately 4 to 13 surrounding whiskers (Ghazanfar and Nicolelis, 1999b); and they are in a position to reliably and temporally integrate these horizontal inputs (Boucsein et al., 2011b). In addition, they have a larger somatic excitation to inhibition ratio (Adesnik and Scanziani, 2010) and receive strikingly fewer inhibitory inputs (Schubert et al., 2007b).

Is the persistent increase in spontaneous activity we observed due to spatiotemporal integration of the structure of the sandpaper sensory input detected by the principal and surrounding whiskers? To test this hypothesis, we trimmed the surrounding whiskers and stimulated only the principal whisker with the same sensory protocol. We recorded MUA using a linear probe spanning the depth of the cortical layers, with each channel of the probe being assigned to a cortical layer by histological and CSD analyses, as described above.

We found that not only did spontaneous activity in layers IV and Vb fail to increase following single-whisker stimulation, but on average neuronal activity in layers II/III, IV, Vb and VI significantly decreased (Fig. 6*A* and *C*). Spontaneous activity in layers II/III decreased by a factor of 0.76±0.05 (P=4.23E-4, N=8) following the first stimulation, and by a factor of 0.67±0.06 (P=0.0044, N=4) following the second stimulation. Layer IV spontaneous activity decreased by a factor of 0.79±0.03 following the first stimulation (P=3.03E-6, N=18, Fig. 6C, empty asterisks) and 0.73±0.03 following the second stimulation (P=3.17E-7, N=15, Fig. 6C, empty asterisks). Layer Vb neuronal activity decreased by a factor of 0.91±0.03 following the first stimulation (P=0.0070, N=15, Fig. 6C, empty asterisks). Layer VI neuronal spontaneous activity decreased by a factor of 0.73±0.06 (P=0.0013, N=7, Fig. 6C, empty asterisks) following the first stimulation. The decrease in layer IV activity was also accompanied by a small but consistent decrease in the strength of the layer IV sink amplitude by a factor of 0.61±0.08 (P=0.0014, N=6) following the first stimulation and a factor of 0.71±0.10 (P=0.0071, N=10) following the second stimulation (Fig. 6B).

**Fig. 6.**
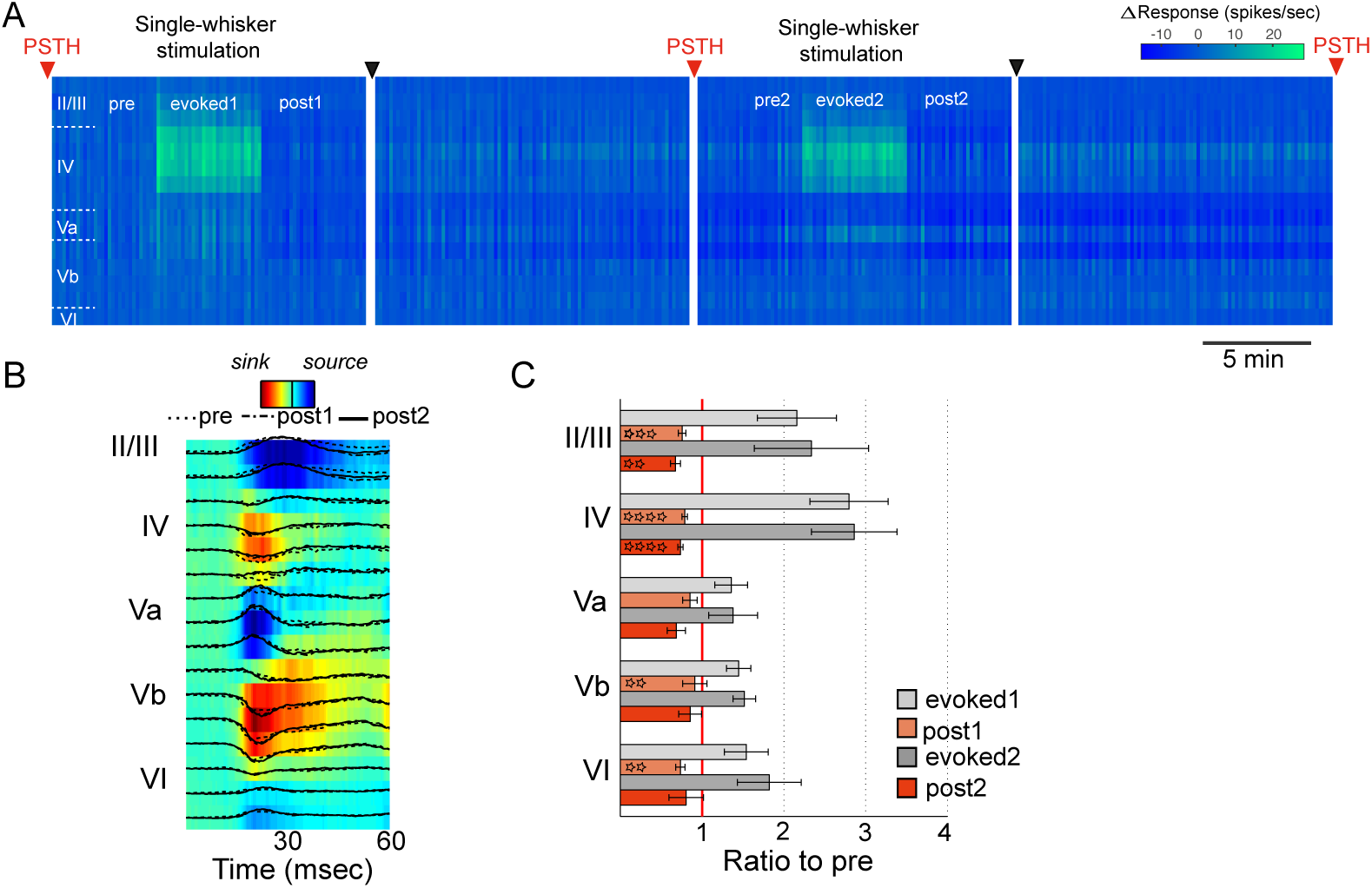
Single-whisker stimulation fails to increase spontaneous activity in layers Vb or IV. (A) Principal whisker stimulation does not increase spontaneous activity in layers IV and Vb: MUA in this example shows failure of spontaneous activity to elevate even following the second sensory stimulation epoch, despite a strong sensory response. (*B*) Single-whisker stimulation leads to a decrease of layer IV sink amplitude (CSD as in Fig. 1, for example shown in panel A). (*C*) Single-whisker stimulation significantly decreases the spontaneous activity in layers II/III, IV, Vb and VI (histograms show changes in firing rate from initial spontaneous levels for channels showing significant sensory responses during the most recent stimulation epoch, as in Fig. 3).

We thus conclude that the increase in spontaneous firing rates in layer IV and Vb requires spatiotemporally rich stimulation across multiple whiskers, which is not present when only the principal whisker is stimulated.

### Sensory patterns of activity reverberate in subsequent spontaneous activity in layers IV and Vb

While the observed multi-unit firing rate increase (Figs. 1-3) is directly related to the neuronal response during stimulation (Fig. *2B*), we further investigated whether the resulting spontaneous activity modifications were due to sensory reverberation. We carried out single-unit recordings to test whether the patterns of activity observed during sensory stimulation reverberate in subsequent spontaneous activity patterns. To achieve this, we used tetrode-configuration probes (2 shanks, Fig. 7*A*) to isolate excitatory single-unit activity (SUA), and repeated the same recording/stimulation protocol.

**Fig. 7.**
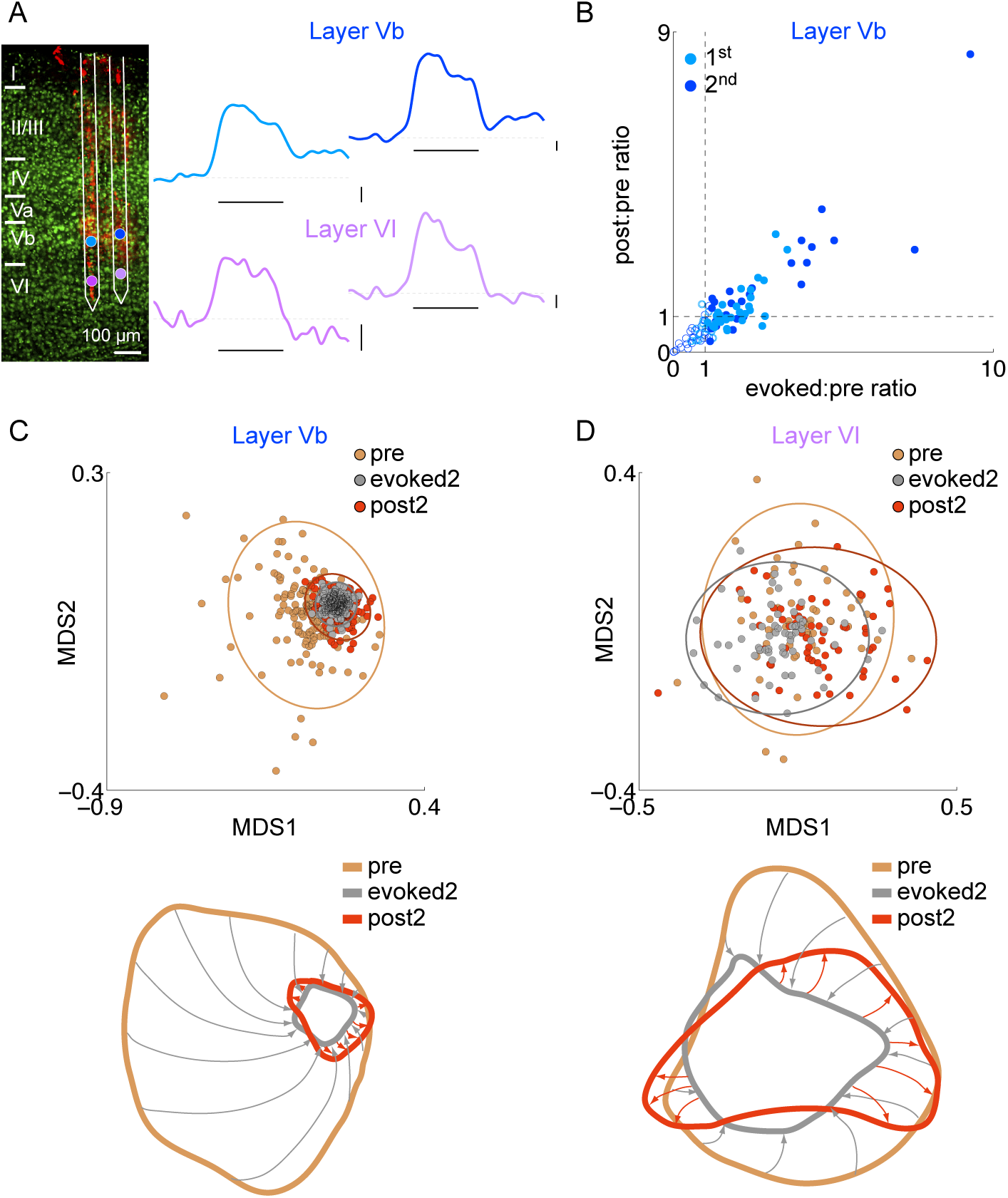
Sensory patterns of activity reverberate into subsequent spontaneous activity. (*A*) Overlap of DiI and Nissl staining of a barrel cortical column showing location of tetrode sites within cortical column. Population firing rates remain at elevated levels following 25 Hz stimulation for tetrodes within layer Vb (blue), but return to baseline for tetrodes within layer VI (purple). Vertical scale bar corresponds to 0.2 spikes/sec; stimulus duration shown by horizontal bar. (*B*) Firing rate increase is directly related to the response during sensory event (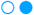 first stimulation, 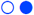 second stimulation). Single units with significant responses are analysed further (filled markers: 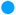 first stimulation, 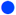 second stimulation). (*C*) MDS was used with the cosine distance to project the vectors of firing rates of single units measured at each tetrode at different times onto a two-dimensional plane (each point corresponds to the state of the group of neurons at a given time-point). The state of co-located neurons converges to a stereotypical pattern of activity during stimulation, as shown by the restricted area in the MDS plane during the evoked epoch. For excitatory neurons in layer Vb, sensory stimulation induces reliable patterns of ensemble activity across time, occupying a smaller volume of MDS space than pre-stimulus spontaneous activity. Post-stimulus spontaneous activity patterns closely resemble the sensory-evoked patterns, as indicated by an overlap integral greater than one between the clouds of points in ‘pre’ and ‘post2’ (*I_r_* = 8.18). (*D*) Sensory stimulation has no effect on post-stimulus spontaneous activity patterns across layer VI single-units and the overlap integral between the cloud of points in ‘pre’ and ‘post2’ is not different from one (*I_r_* = 0.95).

We first examined whether the observed firing rate increases were also detectable in single-units. We found that excitatory SUA in layer Vb increased following 25 Hz sandpaper stimulation and remained elevated thereafter (Fig. 7*A*, blue traces). In contrast, the average SUA firing rate recorded at electrode sites within layer VI returned to baseline following stimulation (Fig. 7*A*, purple traces). In agreement with MUA results, the increase in spontaneous firing of layer Vb excitatory neurons depended strongly on the magnitude of the sensory response (Fig. 7*B*). Layer Vb excitatory neurons with significant sensory responses (Fig. 7*B*, filled markers), increased their spontaneous firing rates by a factor of 1.88±0.35 measured after the second sensory event (p=0.0091, N=23 units).

We next investigated whether the *patterns of activity* produced by the sensory events reverberate in the subsequent spontaneous activity using multidimensional scaling (MDS) to analyse the activity of the neuronal ensemble simultaneously recorded in each experiment. MDS finds a projection for a set of high-dimensional points onto a lower dimensional space that minimally distorts the distances between them, so that data points that are similar in the original higher dimensional space appear close in the projected lower dimensional space (see Materials and Methods). We used MDS to project the time evolution of the state vector of the neuronal ensemble recorded simultaneously in layer Vb and VI onto a 2D space. During the pre-stimulus spontaneous activity, the neuronal ensemble occupies a highly variable pattern space (Fig. 7*C*-*D*, orange) both for layers Vb and VI. In the case of layer VI, the pattern of excitatory single-unit activity converges onto a smaller volume of pattern space during stimulation (compare ‘pre’-orange to ‘evoked2’-grey, Fig. 7*D*), but relaxes back to its original pattern space after the stimulus (compare ‘pre’-orange to ‘post2’-red; Fig. 7*D*). In the case of layer Vb, MDS revealed that the pattern of excitatory single-unit activity converges onto a very small volume of pattern space during sensory stimulation (Fig. 7*C*, ‘evoked2’-grey) indicating a highly similar state during stimulation. Furthermore, the state of layer Vb remains in a similar constrained area of pattern space post-stimulus (Fig. 7*C*, ‘post2’-red). Therefore, in layer Vb, the pattern space occupied post-stimulus resembled that occupied during the sensory event, and both were substantially different to the pattern space occupied prior to stimulation.

To quantify these observations, we obtained the covariance matrices of the points for each epoch (represented by the ellipses), and calculated the *relative overlap integral*, *I_r_*, between the epochs. *I_r_* is a ratio that compares the overlap integral between ‘post2’ and ‘evoked2’ epochs with the overlap integral between ‘pre’ and ‘evoked2’ epochs based on the corresponding clouds of points in the 2D MDS representation (see (Grima et al., 2010) and Materials and Methods). If the states occupy the same space in the MDS representation (i.e. the stimulus has introduced no change in network state), *I_r_* is equal to one. If the ‘post2’ state is more similar to the ‘evoked2’ state than the ‘pre’ state (i.e. the post-stimulus state of the network is more similar to its state during stimulation and considerably different to its pre-stimulus state), then *I_r_* is greater than one. If ‘post2’ is more similar to ‘pre’ than ‘evoked2’ (i.e. the stimulus had an effect, but the network has relaxed back to its original state), then *I_r_* is smaller than one.

The application of this analysis technique can be illustrated with regard to the examples shown in Figure 7 (see also Supplementary Fig. S3). In the case of layer VI, the neuronal pattern state converged during the stimulus, but after stimulation returned to the highly variable pre-stimulus state. The relative overlap integral *I_r_* between ‘pre’ and ‘post2’ was therefore not substantially different from 1 (*I_r_* = 0.95, Fig. 7*D*). In contrast, the patterns in layer Vb converges during the stimulation period, and remain confined during subsequent spontaneous activity, as indicated by a relative overlap integral *I_r_* substantially greater than 1 (*I_r_* = 8.18, Fig. 7*C*). Sufficient recordings with electrode sites in layers IV, Vb and VI were available to calculate population statistics for the overlap integral, pooling 12.5 Hz and 25 Hz stimulation experiments. This behaviour was observed consistently across experiments, with the relative overlap integral being larger than one for layers Vb (*Ī_r_* = 4.18 ± 1.41, mean ± s.d., n=7 animals) and IV (*Ī_r_* = 10.97 ± 5.14, n=3), and not different from one for layer VI (*Ī_r_* = 0.91 ± 0.09, n=3). The reverberation of sensory patterns of activity in subsequent spontaneous activity therefore occurs in layers IV and Vb, but there is not evidence for it occurring in layer VI.

### Fast spiking (FS) interneurons, as well excitatory neurons, show increased spontaneous activity

Putative Fast Spiking (FS) interneurons were identified based on narrow spike waveform characteristics, as described in Methods. Sufficient samples were available to examine FS interneurons in layers IV and Vb (only), with the 12.5 Hz stimulation protocol. Following 12.5 Hz stimulation, units in layers IV and Vb both showed significant elevation of spontaneous activity, similar to that shown following 25 Hz stimulation (see Supplemental Figure S2). We thus were able to examine whether FS interneurons showed the same, or different, dynamics in comparison to broad spiking (putative excitatory) units more common in our recordings, following the sensory stimulation epoch. Table 1 shows the ratio of spontaneous firing rates following to preceding the sensory stimulation epoch; it appears that putative FS interneurons show elevated/reverberant activity in the following epoch, to approximately the same extent as do putative excitatory neurons.

**Table 1.** Separation of units into fast spiking and broad spiking waveforms suggests that putative excitatory and inhibitory units show similar shifts in spontaneous firing rate. n indicates number of units; p the p-value for a one sample Student’s t-test.

|  |  | IV |  |  | Vb |  |  |
| --- | --- | --- | --- | --- | --- | --- | --- |
| **25 Hz stimulation** |  | **Post:pre** | **n** | **p** | **Post:pre** | **n** | **p** |
| Broad spiking |  | Insufficient data | 0 |  | 1.88±0.39 | 30 | 0.0181 |
| Fast spiking |  | Insufficient data | 0 |  | Insufficient data | 2 |  |
| **12.5 Hz stimulation** |  |  |  |  |  |  |  |
| Broad spiking |  | 1.55±0.19 | 37 | 0.088 | 2.32±0.51 | 27 | 0.014 |
| Fast spiking |  | 3.77±1.39 | 8 | 0.0718 | 2.12±0.44 | 14 | 0.02 |

## Discussion

A prominent feature of the somatosensory cortex in rodents is its organization into distinct cortical columns, with each column receiving inputs primarily from one facial whisker and secondary inputs from surrounding whiskers. Rodents rely on spatio-temporally rich information from the collective exploratory motion of their whiskers to identify objects in their environment (Diamond et al., 2008b;Arabzadeh et al., 2005). Multi-whisker stimulation activates lateral circuits via horizontal connections to layers IV (Fox et al., 2003b) and Vb (Ghazanfar and Nicolelis, 1999b), and afferent circuits via multi-whisker thalamocortical projection cells (Bruno and Simons, 2002b). In contrast, single whisker stimulation activates mainly intra-columnar circuits via the corresponding thalamic barreloid. It has been shown that stimulation of adjacent whiskers increases the cross-correlation between neurons in the corresponding cortical columns (Erchova and Diamond, 2004). We therefore reasoned that persistent modification of spontaneous activity might emerge from a naturalistic, spatiotemporally structured stimulus of all whiskers. The present study reveals for the first time persistent, layer-specific modification of spontaneous firing rates by sensory stimulation.

The firing rate increase is specific to layers IV and Vb and lasts for at least 25 minutes poststimulus. Spontaneous firing rate increases following sensory experience have not previously been reported, although potentially related phenomena have been observed, including increases in spontaneous firing rate under a burst-conditioning paradigm (Erchova and Diamond, 2004), reverberation of stimulus-induced voltage-sensitive dye patterns during spontaneous activity (Han et al., 2008), and elevated firing rates in some neurons during the delay period (i.e., after off-set of the initial stimulus) of a working memory task (Fuster and Alexander, 1971;Miller et al., 1993). We speculate that these phenomena are related and that they reflect mechanisms for the maintenance of sensory information in synaptic and/or spontaneous neuronal firing dynamics.

Under Hebbian spike-timing dependent plasticity (STDP) protocols, the synapses are strengthened/depressed depending on the spike-pairing order. This mechanism predicts that patterns of firing repeatedly driven by sensory events should be reflected in subsequent activity due to attractor dynamics (Amit, 1995). In this view, spontaneous activity dynamics reflect underlying circuit structure that is sculpted by the spatiotemporal structure of stimulation throughout a lifetime; in the stimulation protocol employed in the current paper, a substantial amount of spatiotemporally structured stimulation is compressed into a relatively short period of time, leading to exaggerated effects on cortical circuit functional connectivity. Here we show that specifically in layers IV and Vb, the patterns of activity evoked during sensory stimulation in an ensemble of neurons reverberate in subsequent spontaneous activity. In layers IV and Vb, the patterns of activity following sensory events remain substantially different to the highly variable pre-stimulus spontaneous state of activity, resembling more closely instead sensory-evoked activity patterns (Fig. 7). We have previously shown similar results in a computational model of a neuronal network with STDP in the excitatory synapses, where the network converged onto a post-stimulus state sufficiently different from its pre-stimulus state following several repetitions of the same stimulus pattern (Phoka et al., 2012).

Spontaneous activity in layers IV and Vb increases only following multi-whisker stimulation. We surmise that the sustained increase in spontaneous activity in layers IV and Vb therefore depends on the activation of lateral circuits and/or the magnitude of the afferent thalamocortical (or corticothalamocortical) drive, since the effect disappears when mainly intra-columnar circuits are activated (Fig. 6). In support of such a hypothesis involving the modification of the magnitude of the afferent thalamocortical or lateral circuits, is the increase in layer IV amplitude of the CSD profile following the sensory stimulation protocol (Fig. 1). Such changes have previously been related to long-term potentiation (Mégevand et al., 2009). Layers IV and Vb receive direct thalamic (Swadlow et al., 2002) and lateral (Manns et al., 2004;Schubert et al., 2003) drive. Layer IV neurons receive primary horizontal connections from layer IV neurons in surrounding columns (Schubert et al., 2007b), whereas layer Vb neurons receive strong intracolumnar and transcolumnar excitatory inputs from all cortical layers (Schubert et al., 2007b), suggesting the possibility that they integrate (Diamond et al., 2008b;Schubert et al., 2007b;Chagnac-Amitai and Connors, 1989;Boucsein et al., 2011b) these inputs before projecting to downstream brain structures (Alloway, 2008).

Many important questions requiring further research emerge from our findings (Fig. 8). Could blocking the receptors involved in plasticity mechanisms eliminate the firing rate increase? What is the meaning of the differences observed when changing the stimulation frequency? Does the firing rate increase depend on the spatio-temporal structure of the stimulus, and would different sandpaper gratings yield different results? Although much remains to be explored, the possibility that the increases in firing rates are due to spatio-temporal integration remains an attractive hypothesis to explain the observed spontaneous activity modification in response to sensory events. More broadly, one could conjecture that the multi-whisker thalamocortical projections and intracortical horizontal connections play a pivotal role in information integration across multiple facial whiskers. It is therefore of interest to investigate whether similar results could hold in other sensory modalities. The persistent modification of spontaneous dynamics by sensory signals, as demonstrated here, may yield insight into the cortical basis of memory, by elucidating mechanisms for the integration of temporal and spatial context (Diamond et al., 2008b;Schubert et al., 2007b).

**Fig. 8.**
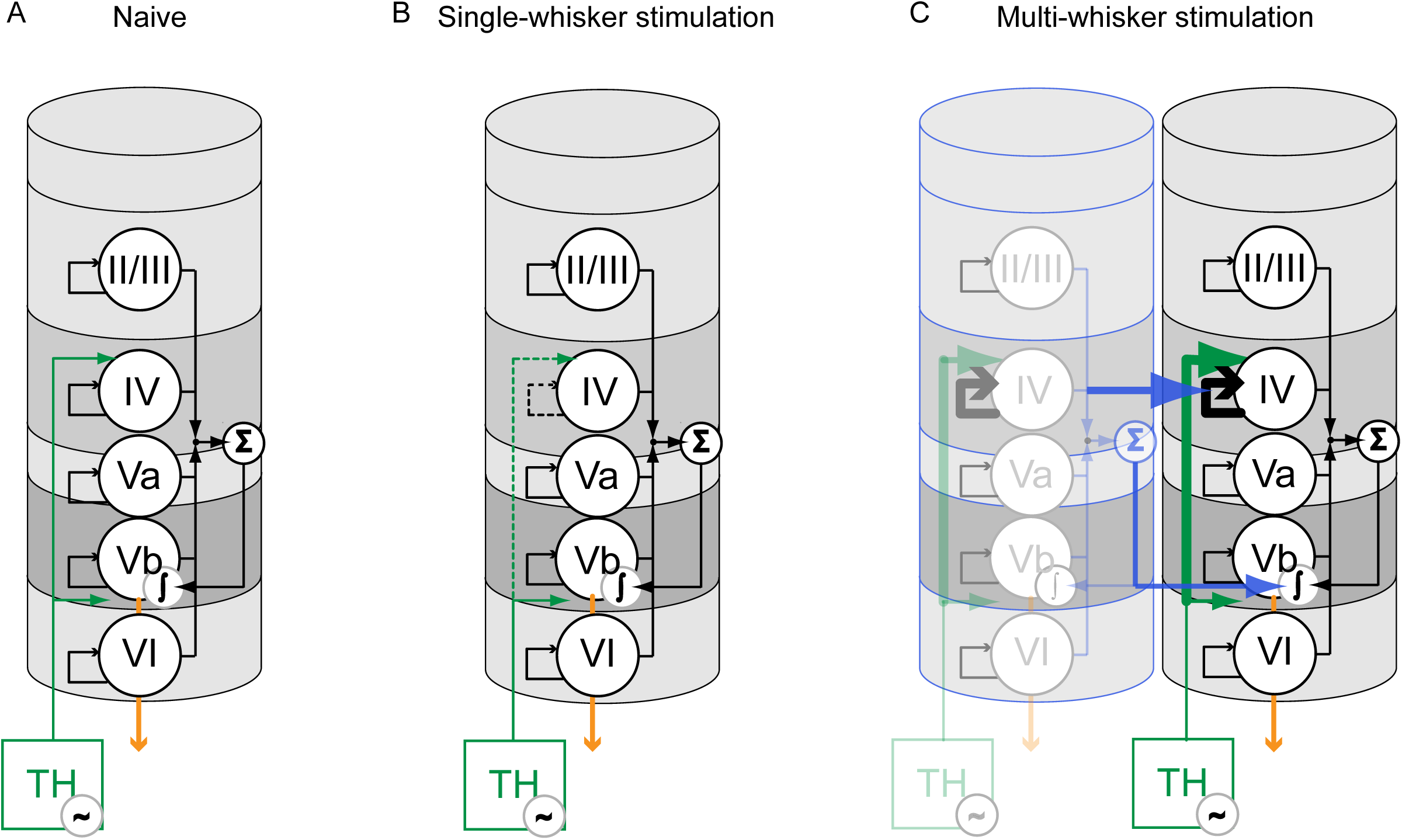
Modification of somatosensory cortical circuitry following multi-whisker stimulation and comparison to single-whisker stimulation. (*A*) A single cortical column consists of five main laminae (II/III, IV, Va, Vb, VI). Cells in each layer receive intra-laminar connections (black arrows). Layers IV and Vb receive thalamocortical inputs (TH, green arrows) from one thalamic barreloid (Fox, 2008). Layer Vb is considered to provide the main output of the cortical column (Schubert et al., 2007a) to downstream subcortical structures (Mercier et al., 1990;Wise and Jones, 1977) (orange arrow). (*B*) Single whisker stimulation mainly activates the corresponding column. We find that stimulation of only one (the primary) whisker leads to a significant decrease in the firing rates in layers II/III, IV, Vb and VI (Fig. 6 *B* and *D*). It also leads to a decrease in the amplitude of the CSD sink amplitude in layer IV (Fig. 6 *C*), which we represent by weakening the thalamocortical and intralayer IV connections (dotted green and black arrows, respectively). (*C*) Multi-whisker stimulation activates the measured barrel column as well as its surrounding columns (represented with a blue outline). We find that multi-whisker stimulation leads to a significant, long-term increase of the activity of layer Vb and IV neurons (Fig. 1 *B*, *E*). This is accompanied by a significant increase in the amplitude of the CSD sink in layer IV (Fig. 1 *C*), represented here by strengthening the thalamocortical, trans-columnar and intra-layer IV connections (bold green, blue and black arrows, respectively). We suggest that multi-whisker stimulation potentiates afferent and/or trans-barrel connectivity via strengthening either thalamocortical (Bruno and Simons, 2002a), or lateral, transcolumnar connections to layers IV (Fox et al., 2003a) and Vb (Ghazanfar and Nicolelis, 1999a). Layer IV neurons receive primary horizontal connections from layer IV neurons from surrounding columns (Schubert et al., 2007a), whereas layer Vb neurons receive strong intracolumnar (black arrows and summation) and transcolumnar excitatory inputs (Schubert et al., 2007a) (blue arrows and summation). This suggests that layer Vb neurons might integrate (Ghazanfar and Nicolelis, 1999a;Boucsein et al., 2011a;Diamond et al., 2008a;Schubert et al., 2007a) multi-whisker sensory inputs before projecting to downstream brain structures (orange arrow).

## Methods

### In vivo physiology

Recordings were made from adult C56BL/6 mice, of 2-3 months of age. All procedures were approved by the UK Home Office under Project License 70/7355 to SR Schultz. For surgery, mice were sedated with an initial intraperitoneal injection of urethane (1.1 g/kg, 10% w/v in saline) followed by an intraperitoneal injection of 1.5 ml/kg of Hypnorm/Hypnovel (a mix of Hypnorm in distilled water 1:1 and Hypnovel in distilled water 1:1; the resulting concentration being Hypnorm:Hypnovel:distilled water at 1:1:2 by volume), 20 minutes later. Atropine (1 ml/Kg, 10% in distilled water) was injected subcutaneously. Further supplements of Hypnorm (1 ml/kg, 10 % in distilled water) were administered intraperitoneally if required.

The mouse's body temperature was maintained at 37±0.5 ºC with a heating pad. A tracheotomy was performed and an endotracheal tube (Hallowell EMC) was inserted to maintain a clear airway as previously described (Moldestad et al., 2009). After the animal was placed on the stereotaxic frame, a craniotomy was performed (approximately 0.5-1 mm in diameter) above barrel C2. A small window in the dura was opened to allow insertion of the multi-electrode array. The exposed cortical surface was covered with artificial cerebrospinal fluid (in mM: 150 NaCl, 2.5 KCl, 10 HEPES, 2 CaCl2, 1 MgCl2; pH 7.3 adjusted with NaOH) to prevent drying. The electrode was lowered into the brain, perpendicularly to the cortical surface, and allowed to settle for 30 minutes before recording began. The eyes were covered with ophthalmic lubricant ointment to prevent drying.

### Data Acquisition and Analysis

Multiunit activity recordings were obtained using a linear probe spanning all cortical layers, with 16 sites spaced at 50 µm intervals (model: A1x16-3mm-50-413, NeuroNexus Technologies). Tetrode probes were used for experiments requiring single-unit isolation (A2x2-tet-3mm-150-312, NeuroNexus Technologies). Signals were acquired with the CED Power1401 data acquisition interface and Spike2 software (Cambridge Electronic Design Limited) and analyzed in Matlab (Mathworks). The extracellular signal was sampled at 20 kHz and local field potential (LFP) signals were extracted by bandpass filtering (1 to 300 Hz). The LFP signals were notch-filtered (centered around 50 Hz) prior to calculation of the power and spectrograms.

For offline spike sorting, the 300 Hz to 9 kHz band was extracted. Spike sorting was performed automatically using KlustaKwik (Harris et al., 2000), followed by manual cluster adjustment using the Klusters software package (Hazan et al., 2006). Units were then classified as fast or broad spiking, based on their peak to trough temporal distance (Niell and Stryker, 2008). Single-units with peak-to-trough distance smaller than 0.4 ms were classified as fast spiking (FS) putative interneurons, whereas units with peak-to-trough distance broader than 0.4 ms were classified as putative excitatory neurons.

### Statistics

To isolate the MUA channels with significant responses (e.g. Fig. 3), we performed a tailed z-test to test the hypothesis that the distribution of the firing rate vector is different to the pre-stimulus firing rate distribution described by its mean and standard deviation. For statistical significance of the firing rate ratios, the one-sample tailed Student’s t-test was used. The statistical tests were performed against the null hypothesis that the mean ratio equals one, at the 95% significance level.

To quantify possible modifications in the CSD sinks (e.g. Fig. 1 *C* and Fig. 5 *C*), we accepted the channels within layer IV and Vb with significant amplitude differences between sink and baseline (within 15 ms of stimulus onset). To do so, we used a Student’s t-test to accept the channels whose mean during baseline was more positive than the sink minimum. We then estimated the statistical significance of the sink amplitude ratios, using one-sample tailed Student’s t-test. The statistical tests were performed against the null hypothesis that the mean ratio is equal to one, at the 95% significance level.

### Stimulation

Rough grade (P40) sandpaper was driven using a servomotor (Futaba S9254) controlled by pulsewidth modulation, swept across the whiskers in the rostro-caudal axis. For multi-whisker stimulation, the whiskers were all cut to the same length of 10 mm immediately prior to electrophysiological recording, following standard procedures (see e.g. Huang et al 1998). For single-whisker stimulation, all whiskers were trimmed to their base except the principal whisker, which was cut to a length of 10 mm. In single-whisker experiments, care was taken to deflect only the principal whisker and not to cause displacement of neighboring whisker follicles. Servomotor characteristics (such as the requirement for a pulse every 20 ms) limited the set of stimulation frequencies that could be applied with this system. For the primary (adapting) stimulus protocol, the servomotor was driven to a squarewave modulated set of positions at 25 or 12.5 Hz, repeated for 5 minutes with a 50% duty cycle (0.8 sec ON, 0.8 sec OFF). Most of the data described in this paper was collected with 25 Hz stimulation, approximately corresponding to the high end of the mouse whisking frequency range.

To obtain Post Stimulus Time Histogram (PSTH) characterization of neural responses, the whiskers were stimulated with brief (80 ms) protractions (-90º) and retractions (+90º) of the sandpaper, with a deflection every 2 seconds (see Supplementary Fig 1). This stimulus was designed such as not to be sufficient to elicit changes in spontaneous activity by itself, while still allowing the system dynamics to be probed. Throughout the paper we refer to this as the PSTH Stimulus. PSTH characterisations using the PSTH stimulus were repeated several times throughout each experiment, both prior to and following the adapting protocol, in order to monitor sensory responsiveness and current source density (CSD) patterns throughout the laminar depth.

### Histology

To determine the position of the electrode in the cortex, histological procedures were performed as previously described (Blanche et al., 2005). Briefly, prior to insertion in the brain, the rear of the electrode shank was painted with orange fluorescent Sulfonated DiI (crystals dissolved in ethanol; SP DiIC18(3), Invitrogen). For histological analysis, animals were overdosed with urethane and transcardially perfused with 1x PBS and 4% formaldehyde (16% formaldehyde in PBS, TAAB Laboratories Equipment Ltd). Brain slices were later treated with green fluorescent Nissl stain (Neurotrace 500/525, Molecular Probes). The electrode track, clearly demarcated by the SP-DiI against the Nissl-stained cortex, was then visualized on a confocal microscope (Leica SP5). Some experiments required the reconstruction of the probe trajectory over multiple sections. The brightness of the red (DiI) channel was adjusted to compensate for the reduced staining with depth of insertion, allowing us to optimize the matching of the electrode sites to anatomical (layer) context. Layer boundaries were assigned on the green (Nissl stained) channel, while the tip of the electrode was determined using the red channel. Merging the two channels then allowed us to determine the layer location of the 16 electrode sites by measuring away from the tip in 50-micron increments (Fig. 1D).

### Current Source Density Analysis (CSD)

To calculate the CSD, a linear probe containing 16 electrode sites with 50 µm spacing (model: A1x16-3mm-50-413, NeuroNexus Technologies) was inserted into the barrel cortex. The signal was filtered (1-300 Hz) and down-sampled to 2 kHz. These local field potentials were averaged across trials with the same stimulus, the mean was subtracted for each electrode, and the MATLAB program *CSDplotter* (Pettersen et al., 2006) was used to calculate the CSD.

### Dimensionality reduction: Multi-Dimensional Scaling (MDS)

In order to study changes of the state of the network due to sensory stimulation, we used MDS as previously described (Phoka et al., 2012). Briefly, MDS operates on a geometric principle: it finds a projection onto a lower dimensional space that distorts minimally the distances between the data points (Kruskal, 1978). As a result, similar measurements in the original dataset remain close in the geometric projection, while dissimilar measurements are kept apart.

Our original state vector consists of the firing rates of N neurons over T time bins (concatenating pre, post2 and evoked2) giving rise to a [N×T] firing rate matrix, *R*. The binning interval was chosen such that at least one neuron is firing over every time bin. The firing rate matrix was used to calculate a [T×T] distance matrix *D*, whose elements correspond to the *cosine distance* between the state vector *rN* ×1 at any two time bins:

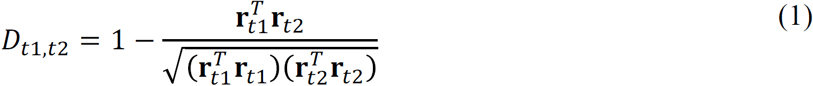

MDS was then used to find a projection of the [N×T] matrix *R* onto 2 dimensions to obtain a [2×T] matrix 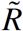 that introduces minimal distortion to *D*.

The cosine distance matrix is used to exclude the possibility that changes in the state vector representation are affected by differences in the absolute magnitude of firing rates. Therefore the representation extracted by MDS represents the similarity of the *patterns* of activity across time, irrespective and exclusive of changes in the average levels of activity.

### Overlap Integral

As a simple measure to quantify the similarity between the cloud of points of the ‘pre’, ‘evoked2’ and ‘post2’ epochs in the 2D MDS representation, we estimate the centroid and covariance matrix of each cloud (represented by the corresponding ellipses) and we calculate their corresponding overlap integrals. Assuming that each ellipse represents a Gaussian probability distribution of the cloud of points, the overlap of two ellipses, *A* and *B*, is given by the overlap integral, *I(A,B)*, as previously described (Grima et al., 2010):

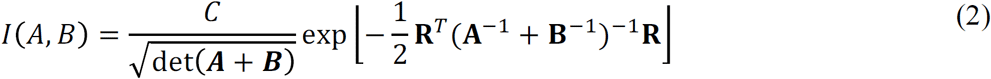

with 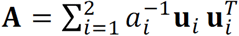 and 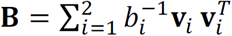, where *a_i_* (respectively, *b_i_*) are the semiaxes along the corresponding normalized eigenvectors **u**_i_ (respectively, **v**_i_) of the ellipses centered at ***R***_*A*_ (respectively, ***R***_*B*_), as obtained by diagonalizing the estimated covariance matrices. 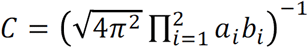 normalization constant and **R**=**R**_B_-**R**_A_ is the distance vector between the two ellipses.

This calculation is equivalent to obtaining the (undisplaced) convolution of two anisotropic Gaussians centered at different points of the MDS 2D space. Therefore, the overlap integral gives a measure of the intersection of the two distributions relative to their own variances.

To quantify the difference between ‘pre’ and ‘post2’ states, we define the *relative overlap integral* as the ratio of the (post2,evoked2) overlap integral over the (pre,evoked2) overlap integral:

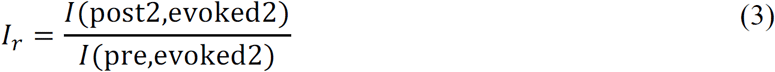

If the ‘pre’ and ‘post2’ states occupy the same position in the MDS representation, then *I_r_* = 1. If *I_r_ > 1*, then the ‘post2’ state has more in common with the ‘evoked2’ state than ‘pre’ does. If *I_r_ < 1*, then ‘pre’ is more similar to ‘evoked2’ than ‘post2’. The values of *I_r_* have been calculated across experiments to quantify the similarity of the ‘pre’, ‘evoked2’ and ‘post2’ for neuronal ensembles in different layers.

## Acknowledgments

We thank G. Hadjithomas for the drawings; A. Saleem and T. Litke for technical assistance; M. Diamond, K. Harris, K.H. Parker and D. Schubert for critical discussions. This work was supported by BBSRC DTA studentships (E.P. and A.B.), and grants from the BBSRC BB/K001817/1 (SRS), and the EPSRC EP/I017267/1 (M.B.).

The authors declare no conflict of interest.

## Figure Legends

**Fig. S1.**
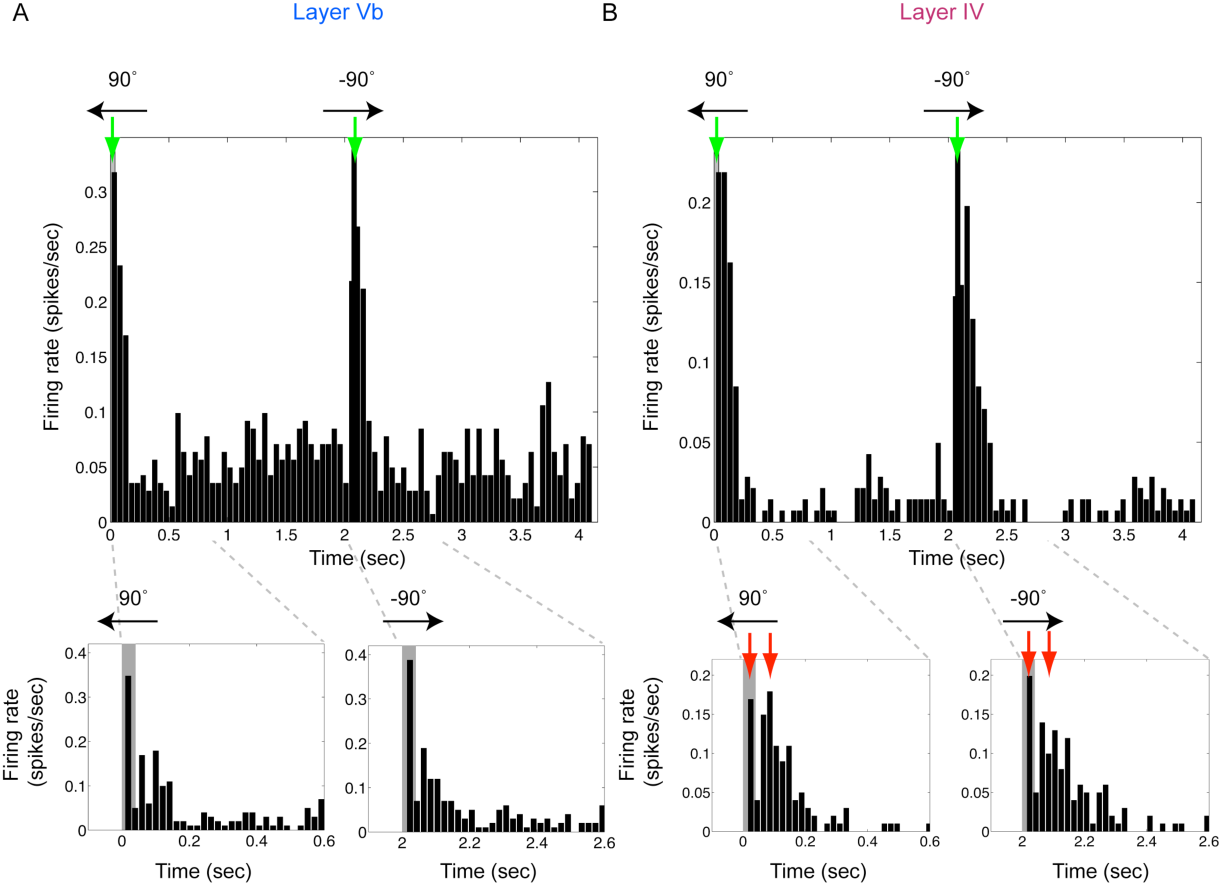
Reliable and temporally precise responses to brief stimulation with sandpaper demonstrated by post-stimulus time histograms (PSTHs) of MUA for the step stimulus. In order to evaluate how well the neurons responded to the stimulus, and in order to identify putative direct thalamocortical input locations within the cortical column using CSD analysis, we stimulated the whiskers with brief (80 ms) deflections using sandpaper. The sandpaper was swept across the vibrissae every 2 seconds in mirrored protraction (-90°) and retraction (90°) directions. This protocol was performed at the very start of the recordings, in the middle of the two stimulation epochs and at the very end of the recordings (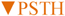, Fig. 1). PSTHs shown here were obtained with 25 Hz stimulation from: (*A*) layer Vb and (*B*) layer IV. Neuronal responses align to stimulus onset (upper panels, green arrows). PSTHs on shorter time scales (lower panels) show two peaks (red arrows) attributed to a characteristic centresurround triphasic component (1) (early depolarization-hyperpolarization-rebound depolarization). The LFP component of the recorded signal during these brief stimulations was used for the CSD analysis.

**Figure S2.**
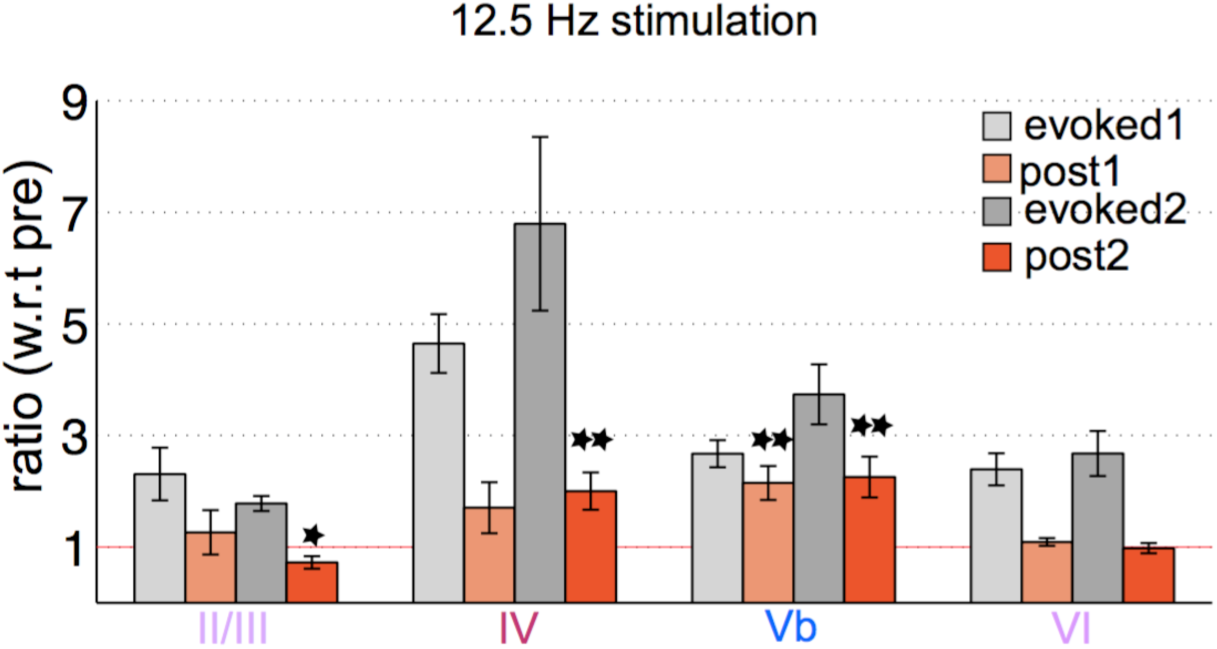
Stimulation at 12.5 Hz results in broadly similar, layer-specific shifts in spontaneous SUA, to that observed with stimulation at 25 Hz, and to that observed with MUA. Bar charts show the ratio between the average firing rate over the labelled five-minute epoch, to that during the five minutes prior to onset of the stimulation epoch. Error bars indicate standard error of the mean over units isolated from each of layer (n=6 animals, 15/11, 39/40, 64/40, 21/17 units respectively, with inclusion criterion of responding significantly during the most recent sensory stimulation epoch, for layers II/III through VI respectively). Single and double asterisks indicate statistical significance at p<0.05 and p<0.01 respectively (Student’s t-test).

**Fig. S3.**
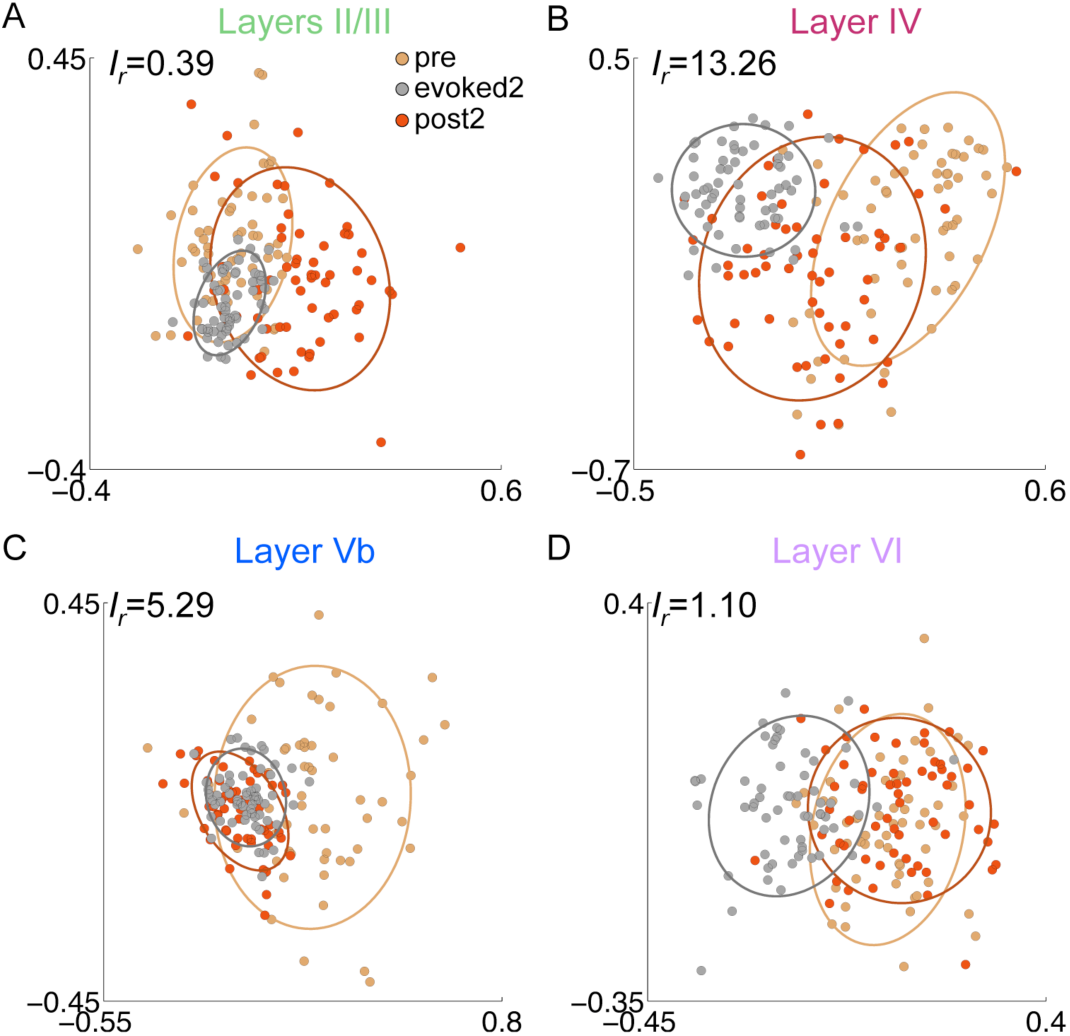
Multi-dimensional Scaling (MDS) analysis of the SUA data reveals distinct changes in the firing patterns of single units in different layers, following 12.5 Hz as well as 25 Hz stimulation (see Fig. 7 C and *D* for 25 Hz results). In layers IV and Vb, the similarity between the post-stimulus and evoked states is higher than between the pre-stimulus and evoked states. This effect is apparently not present in layers II/III and VI. This figure shows an example; overlap integral statistics summarizing the population of recordings across animals are provided in the Results section. (A) Layer II/III neurons have an evoked2 state more similar to the pre-stimulus rather than post-stimulus state (*I*_*r*_ = 0.39). (B) The post-stimulus state of neurons in layer IV is highly similar to evoked2 and distinct from pre (*I*_*r*_ = 13.26). (C) Starting from a large region in the pre-stimulus state, the state of the neurons in layer Vb becomes confined to a small area of the plane during stimulation and remains so post-stimulus (*I*_*r*_ = 5.29). (D) The post- and pre-stimulus states of neurons in layer VI are similar to each other and dissimilar to the evoked2 state (*I*_*r*_ = 1.10).

## References

Adesnik H, Scanziani M (2010) Lateral competition for cortical space by layer-specific horizontal inputs. Nature 464:1155–1160.

Alloway KD (2008) Information processing streams in rodent barrel cortex: the differential functions of barrel and septal circuits. Cereb Cortex 18:979–989.

Amit DJ (1995) The hebbian paradigm reintegrated: local reverberations as internal representation. Behav Brain Sci 18:617–626.

Arabzadeh E, Zorzin E, Diamond ME (2005) Neuronal encoding of texture in the whisker sensory pathway. PLoS Biol 3:e17.

Arieli A, Sterkin A, Grinvald A, Aersten A (1996) Dynamics of ongoing activity: explanation of the large variability in evoked cortical responses. Science 273:1868–1871.

Azouz R, Gray C (1999) Cellular mechanism contributing to response variability of cortical neurons in vivo. J Neurosci 19:2209–2223.

Blanche TJ, Spacek MA, Hetke JF, Swindale NV (2005) Polytrodes: high-density silicon electrode arrays for large-scale multiunit recording. J Neurophysiol 93:2987–3000.

Boucsein C, Nawrot MP, Schnepel P, Aersten A (2011) Beyond the cortical column: abundance and physiology of horizontal connections imply a strong role for inputs from the surround. Front Neurosci 5:1–13.

Brecht M, Grinevich V, Jin TE, Margrie T, Osten P (2006) Cellular mechanisms of motor control in the vibrissal system. Pflügers Arch. Eur. J. Physiol. 453:269–281.

Bruno RM, Simons DJ (2002) Feedforward mechanisms of excitatory and inhibitory receptive fields. J Neurosci 22:10966–10975.

Chagnac-Amitai Y, Connors BW (1989) Synchronized excitation and inhibition driven by intrinsically bursting neurons in neocortex. J Neurophysiol 62:1149–1162.

Chauvette S, Crochet S, Volgushev M, Timofeev I (2011) Properties of slow oscillation during slow-wave sleep and anesthesia in cats. J Neurosci 31:14998–15009.

Diamond ME, von Heimendahl M, Knutsen PM, Kleinfeld D, Ahissar E (2008) ‘Where’ and ‘what’ in the whisker sensorimotor system. Nature Rev Neurosci 9:601–612.

Erchova IA, Diamond ME (2004) Rapid fluctuations in rat barrel cortex plasticity. J Neurosci 24:5931–5941.

Erchova IA, Lebedev MA, Diamond ME (2002) Somatosensory cortical neuronal population activity across states of anaesthesia. European Journal of Neuroscience 15:744–752.

Ferezou I, Haiss F, Gentet LJ, Aronoff R, Weber B, Petersen CCH (2007) Spatiotemporal dynamics of cortical sensorimotor integration in behaving mice. Neuron 56:907–923.

Fox K (2008) Barrel Cortex. Cambridge, UK: Cambridge University Press.

Fox K, Wright N, Wallace H, Glazewski S (2003) The origin of cortical surround receptive fields studied in the barrel cortex. J Neurosci 23:8380–8391.

Fox MD, Raichle ME (2007) Spontaneous fluctuations in brain activity observed with functional magnetic resonance imaging. Nat Rev Neurosci 8:700–711.

Freeman JA, Nicholson C (1975) Experimental optimization of current-source density technique for anuran cerebellum. J. Neurophysiol. 38:369–382.

Fuster JM, Alexander GE (1971) Neuron activity related to short-term memory. Science 173:652–654.

Ghazanfar AA, Nicolelis MA (1999) Spatiotemporal properties of layer V neurons of the rat primary somatosensory cortex. Cereb Cortex 9:348–361.

Goard M, Dan Y (2009) Basal forebrain activation enhances cortical coding of natural scenes. Nature Neurosci 12:1444–1449.

Goldman-Rakic PS (1995) Cellular basis of working memory. Neuron 14:477–485.

Grima R, Yaliraki SN, Barahona M (2010) Crowding-induced anisotropic transport modulates reaction kinetics in nanoscale porous media. J Phys Chem B 114:5380–5385.

Han F, Caporale N, Dan Y (2008) Reverberation of recent visual experience in spontaneous cortical waves. Neuron 60:321–327.

Harris KD, Henze DA, Csicsvari J, Hirase H, Buzsáki G (2000) Accuracy of tetrode spike separation as determined by simultaneous intracellular and extracellular measurements. Journal of Neurophysiology 84:401–414.

Harris KD, Thiele A (2011) Cortical state and attention. Nature Rev. Neurosci. 12:509–523.

Hazan L, Zugaro M, Buzsáki G (2006) Klusters, NeuroScope, NDManager: a free software suite for neurophysiological data processing and visualization. Journal of neuroscience methods 155:207–216.

Huang W, Armstrong-James M, Rema V, Diamond ME and Ebner FF (1998). Contribution of supragranular layers to sensory processing and plasticity in adult rat barrel cortex. J Neurophysiol 80:3261–71.

Jadhav SP, Wolfe J, Feldman DE (2009) Sparse temporal coding of elementary tactile features during active whisker sensation. Nat Neurosci 12:792–800.

Jin TE, Witzemann V, Brecht M (2004) Fiber types of the intrinsic whisker muscle and whisking behavior. J Neurosci 24:3386–3393.

Kenet T, Bibitchkov D, Tsodyks M, Grinvald A, Arieli A (2003) Spontaneously emerging cortical representations of visual attributes. Nature 425:954–956.

Kirkby LA, Sack GS, Firl A, Feller MB (2014) A Role for Correlated Spontaneous Activity in the Assembly of Neural Circuits. Neuron 80:1129–44.

Knott GW, Quairiaux C, Genoud C, Welker E (2002) Formation of dendritic spines with GABAergic synapses induced by whisker stimulation in adult mice. Neuron 34:265–273.

Kruskal JB (1978) Multidimensional Scaling. SAGE Publications.

Lewis CM, Baldassarre A, Committeri G, Romani GL, Corbetta M (2009) Learning sculpts the spontaneous activity of the resting human brain. Proc Natl Acad Sci U S A 106:17558–17563.

Lottem E, Azouz R (2009) Mechanisms of tactile information transmission through whisker vibrations. J Neurosci 29:11686–11697.

Manns ID, Sakmann B, Brecht M (2004) Sub- and suprathreshold receptive field properties of pyramidal neurones in layers 5A and 5B of rat somatosensory barrel cortex. J Physiol 556:601–622.

Marguet SL, Harris KD (2011) State-dependent representation of amplitude-modulated noise stimuli in rat auditory cortex. J Neurosci 31:6414–6420.

Mercier BE, Legg CR, Glickstein M (1990) Basal ganglia and cerebellum receive different somatosensory information in rats. Proceedings of the National Academy of Sciences 87:4388–4392.

Mégevand P, Troncoso E, Quariaux C, Muller M, Michel MC, Kiss ZJ (2009) Long-term plasticity in mouse sensorimotor circuits after rhythmic whisker stimulation. J Neurosci 29:5326–5335.

Miller EK, Li L, Desimone R (1993) Activity of neurons in anterior inferior temporal cortex during a short-term memory task. J Neurosci 13:1460–1478.

Mitra A and Raichle ME (2016). How networks communicate: propagation patterns in spontaneous brain activity. Phil. Trans. R. Soc. B 371: 20150546

Mitzdorf U (1985) Current source-density method and application in cat cerebral cortex: investigration of evoked potentials and EEG phenomena. Physiol Rev 65:37–100.

Moldestad O, Karlsen P, Molden S, Storm JF (2009) Tracheotomy improves experiment success rate in mice during urethane anesthesia and stereotaxic surgery. Journal of neuroscience methods 176:57–62.

Niell CM, Stryker MP (2008) Highly selective receptive fields in mouse visual cortex. J Neurosci 28:7520–7536.

Pettersen KH, Devor A, Ulbert I, Dale AM, Einevoll GT (2006) Current-source density estimation based on inversion of electrostatic forward solution: Effects of finite extent of neuronal activity and conductivity discontinuities. J Neurosci Methods 154:116–133.

Phoka E, Wildie M, Schultz SR, Barahona M (2012) Sensory experience modifies spontaneous state dynamics in a large-scale barrel cortical model. J Comput Neurosci, 33:323–339.

Quairiaux C, Armstrong-James M, Welker E (2007) Modified sensory processing in the barrel cortex of the adult mouse after chronic whisker stimulation. J Neurophysiol 97:2130–2147.

Reyes-Puerta R, Yang J-W, Siwek ME, Kilb W, Sun J-J, Luhman HJ (2016). Propagation of spontaneous slow-wave activity across columns and layers of the adult rat barrel cortex in vivo. Brain Structure and Function, doi:10.1007/s00429-015-1173-x.

Ringach DL (2009) Spontaneous and driven cortical activity: implications for computation. Curr Opin Neurobiol 19:439–444.

Schubert D, Kötter R, Staiger JF (2007a) Mapping functional connectivity in barrel-related columns reveals layer- and cell type-specific microcircuits. Brain Struct Funct 212:107–119.

Schubert D, Kötter R, Staiger JF (2007b) Mapping functional connectivity in barrel-related columns reveals layer- and cell type-specific microcircuits. Brain Struct Funct 212:107–119.

Schubert D, Kötter R, Zilles K, Luhmann HJ, Staiger JF (2003) Cell type-specific circuits of cortical layer IV spiny neurons. J Neurosci 23:2961–2970.

Sokoloff L, Mangold R, Wechsler RL, Kenney C, Kety SS (1955) The effect of mental arithmetic on cerebral circulation and metabolism. J Clin Invest 34:1101–1108.

Swadlow HA, Gusev AG, Bezdudnaya T (2002) Activation of a cortical column by a thalamocortical impulse. J Neurosci 22:7766–7773.

Tegnér J, Compte A, Wang X-J (2002) The dynamical stability of reverberatory neural circuits. Biol Cybern 87:471–481.

Tsodyks M, Kenet T, Grinvald A, Arieli A (1999) Linking spontaneous activity of single cortical neurons and the underlying functional architecture. Science 286:1943–1946.

Wise SP, Jones EG (1977) Somatotopic and columnar organization in the corticotectal projection of the rat somatic sensory cortex. Brain research 133:223–235.

Yao H, Shi L, Han F, Gao H, Dan Y (2007) Rapid learning in cortical coding of visual scenes. Nat Neurosci 10:772–778.

